# Sequestration of dead-end undecaprenyl phosphate-linked oligosaccharide intermediate

**DOI:** 10.1101/2024.09.22.614228

**Authors:** Yaoqin Hong, Peter R. Reeves

## Abstract

In the predominant Wzx/Wzy-dependent bacterial surface polysaccharide biosynthetic pathway, synthesis is divided between the cytoplasmic and periplasmic faces of the membrane. Initially, an oligosaccharide composed of 3-8 sugars is synthesized on a membrane-embedded lipid carrier, Und-P, within the cytoplasmic face of the membrane. This Und-P-linked oligosaccharide is then translocated to the periplasmic face by the Wzx flippase, where it is polymerized into a repeat-unit polysaccharide. Structural alterations to the repeating oligosaccharide significantly reduce polysaccharide yield and lead to cell death or morphological abnormalities. These effects are attributed to the substrate recognition function of the Wzx flippase, which we postulated to act as a gatekeeper to ensure only complete substrates are translocated to the periplasmic face. Here, we labelled *Salmonella enterica* serovar Typhimurium Group B1 with [^14^C] D-galactose. Our results showed that strains unable to synthesize the full O-antigen repeat unit accumulate significantly higher levels of Und-P-linked material (∼10-fold). Importantly, this sequestration is alleviated by mild membrane disruption which opens the cytosolic face Und-PP-linked material to O-antigen ligation that supports the accumulation to occur at the cytosolic face of the membrane.

## INTRODUCTION

The synthesis of glycoconjugates on lipid carriers is a conserved anabolic process across all domains of life. In bacteria (excluding mycobacteria), undecaprenyl phosphate (Und-P) is the predominant lipid carrier involved in glycan and proteoglycan biosynthesis. This common carrier plays a crucial role in various pathways including peptidoglycan, capsular polysaccharides, exopolysaccharides (e.g., colanic acid), Enterobacteriaceae common antigens (ECA), and O antigen in Enterobacteriaceae species (1). Many bacterial surface polysaccharides are synthesised by the Wzx/Wzy-dependent pathway. In this system, the biosynthetic process begins with the addition of a sugar-phosphate to Und-P by an initial transferase (IT), forming an Und-PP-linked sugar. The oligosaccharide repeating unit is then assembled sequentially and translocated to the periplasm for polymerization by the Wzy polymerase (2-6). Subsequently, the released Und-PP is dephosphorylated and recycled back to the cytoplasmic face of the membrane (7,8).

Relative to the well-established Gram-positive model species, the availability of Und-P is limited in the Gram-negative bacteria (9). Mutations in surface polysaccharide biosynthesis genes often lead to cell death or severe phenotypic consequences, including compromised cell envelope integrity (10-21). One plausible explanation for these effects is that sequestration of the Und-P pool impairs the ability to meet the demands of various biosynthetic pathways, particularly peptidoglycan biogenesis. Indeed, the overproduction of Und-P has been shown to alleviate cell envelope defects, underscoring its crucial role (21). Thus, the traffic in the competing pathways in which Und-P is shared is anticipated to be well managed, and this element has captivated interest in unravelling the underlying interactions governing Und-P flow.

Despite the importance of directly assessing Und-P levels, it has been challenging due to its relatively low abundance compared to other membrane lipids (22). Indeed, the quantitative evidence for Und-P sequestration has primarily come from studies of *Escherichia coli* mutants with truncated ECA oligosaccharide repeat units (23). Previous research has also demonstrated that strains producing truncated O-antigen repeat units exhibit minimal O-antigen production and have severe growth defects (13-15). To mitigate these effects, tactics involving controlled O-antigen production have been employed (13-15). Similarly, Nikaido and colleagues had observed in 1969 that mutants lacking biosynthetic control were unstable and were replaced by revertants or suppressors after overnight growth (10). In our strains in which O-antigen over-production can be switched on, the consequent rapid cell death is attributed to substrate preference at the membrane translocation step, leading to the accumulation of Und-PP-linked material on the cytoplasmic face of the membrane and preventing effective recycling (14,15,24). In this study, we employed radioactive sugar labelling to investigate Und-P sequestration in O-antigen-defective mutants, which provided support for the accumulation occurring on the cytosolic face of the inner membrane.

## RESULTS

### Sequestration of undecaprenyl phosphate in a group B1 O-antigen repeat unit mutant

D-galactose (D-Gal) is the first sugar of the O-antigen repeat unit in *S. enterica* Sv. Typhimurium (Group B1) and an essential component of the outer core of lipopolysaccharide (Fig 1). We used a previously defined Δ*galE* mutant in this work, for which the only source for the acquisition of UDP-Gal is through the GalK/GalT salvage pathway (Supplemental material, Fig S1A). We confirmed that the strain is unable to metabolise D-Gal (See Material and Methods; Supplemental material, Fig S1B), thus allowing dual-labelling of the O antigen and the core oligosaccharide. For simplicity, we will treat the Δ*galE* strain P9528 as our WT strain from here on.

**Fig 1.**
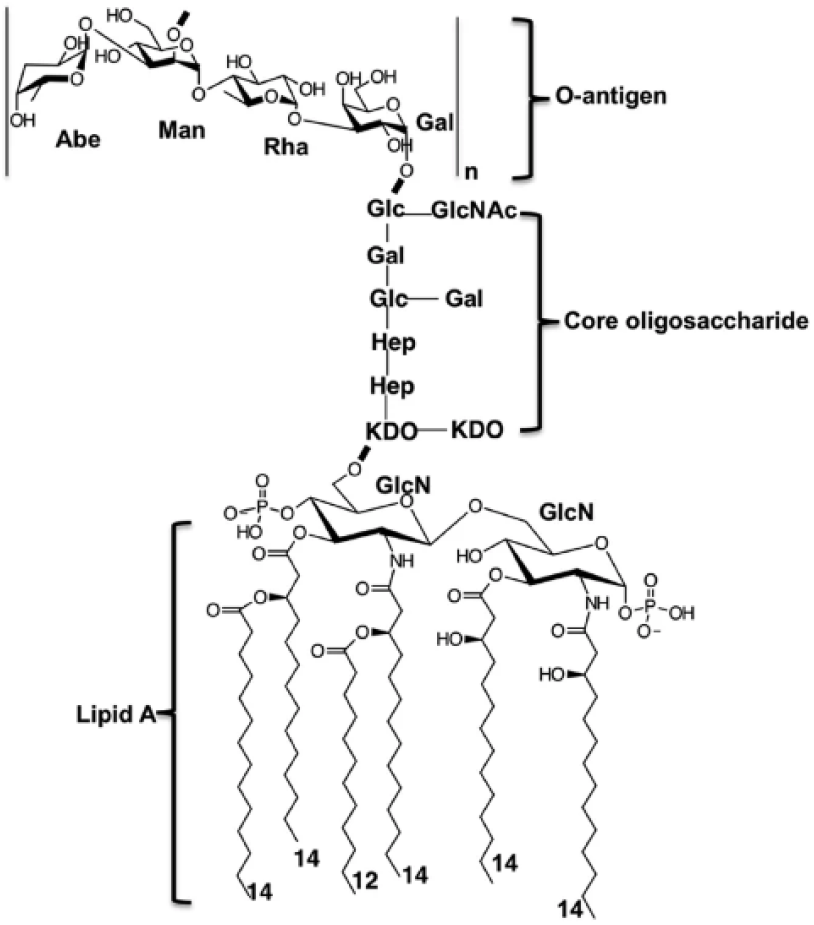
The structure of *S. enterica* Group B1 LPS. Abbreviations: Abe, abequose; Man, mannose; Rha, rhamnose; Gal, galactose; GlcN, glucosamine.

The O-antigen repeat unit of Group B1 *Salmonella* consists of a three-sugar mainchain (mannosyl-rhamnosyl-galactose) with an abequose side branch attached to the mannosyl residue (Fig 1). The *abe* null mutant is unable to synthesise CDP-abequose, and such strains produce only the mainchain structure. Mutants carrying this defect were among the first strains to show lethality after the start of the O-antigen production (10). As reported in our earlier studies (13), the LPS produced by the mutant carries minimal amounts of polymeric O antigen. Mid-log strain cultures with adjusted equivalent OD_600_ (see Material and Methods) were fed with [^14^C] D-Gal for 15 min. For the WT strain, 45% of the[^14^C] D-Gal was taken up by cells. In marked contrast, the uptake efficacy for the Δ*abe* strain for [^14^C] D-Gal (∼15%), was 3-fold less relative to the WT (Fig 2A). Given the majority of [^14^C] D-Gal was retained in the media for both tested strains, the amount of [^14^C] D-Gal, the specific activity, and the uptake duration were considered suitable and used for subsequent assessments.

**Fig 2.**
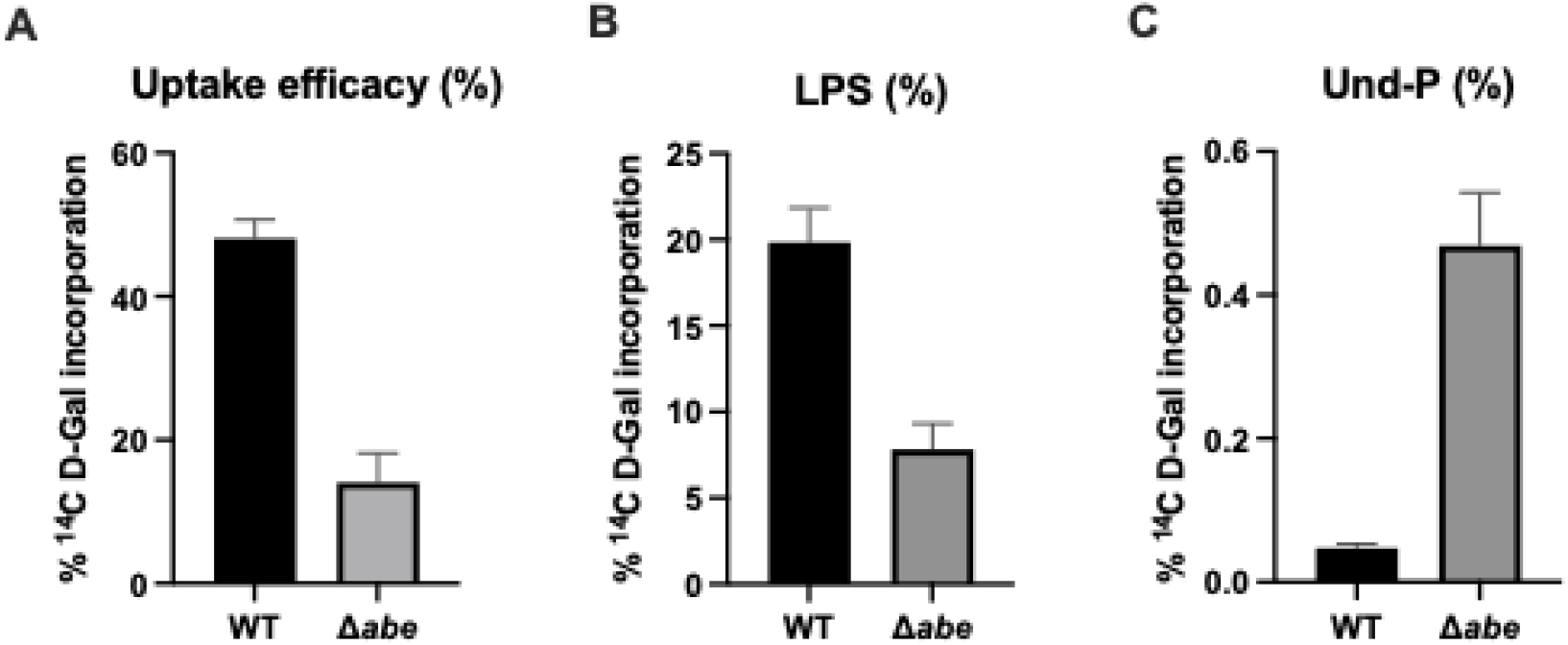
[^14^C] D-Gal uptake in *S. enterica* Group B1 WT and Δ*abe*. A, % [^14^C] D-Gal counts of the whole cell; B, % [^14^C] D-Gal counts in purified LPS; C, % [^14^C] D-Gal counts in n-butanol extracts. Data from three independent repeats are shown and the error bar uses SEM.

Cells from WT and Δ*abe* strains, labelled with [^14^C] D-Gal under the described conditions, were used to quantify the uptake of [^14^C] Gal into LPS. We observed that the [^14^C] D-Gal content incorporated into the phenol-extracted LPS phase of the WT strain was approximately 3-fold higher than that of the Δ*abe* strain (Fig 2B). This finding aligns with our previous observations that the LPS produced by the Δ*abe* strain contains very little O antigen and is mostly comprised of short-chain molecules (13,15). We attributed this to most of the [^14^C] D-Gal being incorporated into the LPS outer core (Fig 1A).

Previously, we suggested that the truncated O-antigen intermediate produced by the Δ*abe* strain is sequestered at the surface translocation step (13,15). To confirm this, we extracted and quantified the Und-PP-linked O-antigen intermediate using n-butanol extraction. The results showed that the [^14^C] D-Gal counts in the n-butanol extract of the Δ*abe* strain were 10-fold higher than those in the WT extract (Fig 2C). This suggests that the WT counts represent the regular Und-P pool allocated to the intended pathway, while the ten-fold higher counts in the n-butanol extract of the Δ*abe* mutant indicate that a significant portion of lipid carriers are sequestered during the 15-minute uptake period. We attributed the high n-butanol counts to an ineffective Wzx flippase.

### Undecaprenyl phosphate sequestration occurred at the cytosolic face of the cytoplasmic membrane

The sequestered Und-PP-linked oligosaccharides were most likely trapped on the cytosolic face of the membrane for several reasons. Firstly, there is substantial genetic and biochemical evidence that the highly variable Wzx proteins function as the cognate repeat-unit flippase (25,26) and each functions only for an appropriate variant or several variants (13,27-30). Secondly, ectopically overexpressing the wzx gene or switching to an appropriate Wzx can reverse the growth defect and repair the O-antigen biosynthesis pathway (13,14,24). To validate this further, we rearranged the cell envelope with repeated mild sonication. This method creates segments of the cytoplasmic membrane with materials from one face of the membrane being adjacent to materials from the opposite face (31). Such rearrangement would facilitate the release of sequestered oligosaccharides through WaaL ligase activity, leading to a significant decrease in [^14^C] D-Gal counts in the n-butanol extract.

Samples were subjected to repeated sonication, and changes in [^14^C] D-Gal counts in the purified LPS and n-butanol extractable fractions were assessed. The sonication resulted in a 43% reduction in [^14^C] D-Gal signals in the n-butanol extract from the Δ*abe* strain compared to the non-sonicated control (Paired *t*-test, P=0.0098) (Fig. 3B). In contrast, there was only a minor change in the [^14^C] D-Gal counts in the n-butanol extract from the WT strain after sonication (Paired *t*-test, P=0.5228) (Fig. 3A). These results strongly suggest that the sequestration of dead-end intermediates occurs specifically at the cytosolic face of the cell membrane.

**Fig 3.**
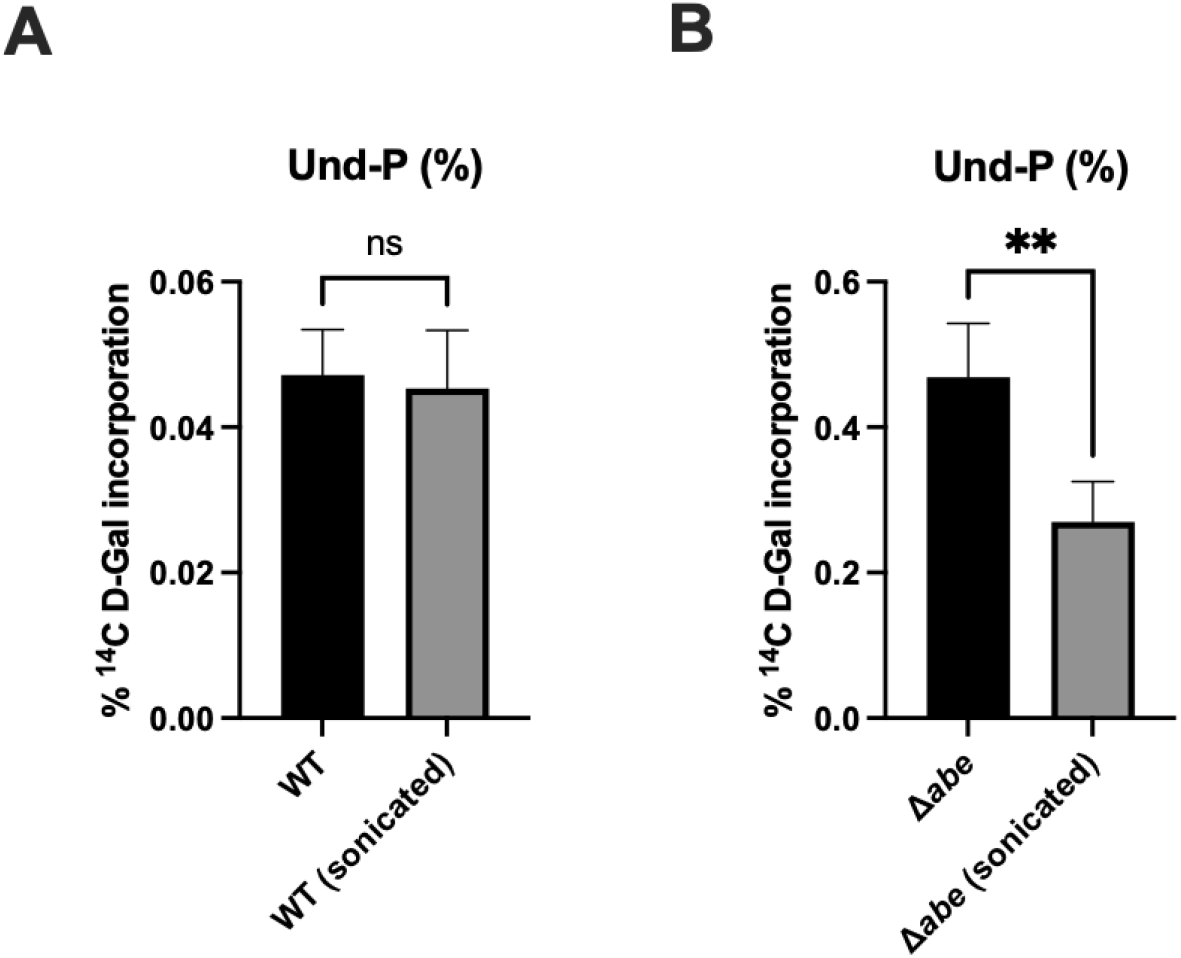
Reduction of [^14^C] D-Gal counts by membrane rearrangements in the n-butanol extractable pool from *S. enterica* Group B1 Δ*abe* but not that from the WT pool. A, Sonication does not affect the n-butanol extractable pool in the WT; B, Sonication significantly reduced the n-butanol extractable pool in the Δ*abe* mutant. Note that the unsonicated n-butanol extractable pool counts in Fig 3A and 3B were the same datasets as Fig 2C. Data from three independent repeats are shown and the error bar uses SEM.

## DISCUSSION

The Und-P lipid carrier is crucial for the biosynthesis of peptidoglycan and surface polysaccharides in most bacterial species. The WecA-initiation pathway is a well-characterized example that is prevalent in *E. coli* (including *Shigella*) and *S. enterica*, where the Und-PP-linked *N*-acetylglucosamine (GlcNAc) product can be channelled into ECA or O-antigen production (32,33). It has long been envisaged that the GTs would constitute a multi-protein complex. Under normal conditions, the pool of lipid carriers is likely regulated by the activity of specific glycosyltransferases (GTs) at the committed steps. In the GlcNAc-initiated pathway, the direction of synthesis is also complicated by the ratio of WecA recruited to respective pathways. The use of group B O:4 in this work evaded this complication to allow a direct assessment of Und-P sequestration.

Defects in Und-P utilisation typically result in poor growth, cell lysis, and morphological abnormalities (10-17,19-21). Previous studies have highlighted the role of Wzx translocase substrate preference in understanding Und-P sequestration and its detrimental effects on cells (13-15,18,21,24). However, direct assessment of Und-P sequestration has been challenging. The only direct evidence available is LC-MS data from methanol-chloroform-water extracted lipids in mutants of the ECA biosynthetic pathway (23). Here, we utilized strains with defects in O-antigen repeat unit structure to capture Und-PP-linked O-antigen intermediates and assessed their sequestration.

Osborn and co-workers assayed the formation of Und-PP-linked O-antigen polymers using radioactively labelled nucleotide sugars with repeatedly sonicated cells to assemble into Und-PP-linked repeat-unit polymer. In this case, the membrane must have been sufficiently rearranged to bring the initial GT and Wzy polymerase to the same membrane leaflet for the reaction to occur (34). We leveraged the capacity of this approach to alleviate stalled Und-PP-linked intermediate and demonstrated that mild sonication significantly reduced the pool of sequestrated Und-PP-linked O-units, indicating that these intermediates accumulate on the cytosolic face of the membrane. The conversion of the Und-PP-linked oligosaccharide into the LPS after sonication appeared insignificant in both the WT and Δ*abe* strains (Supplemental material, Fig S2A). However, we anticipated this would be the case, given the LPS fraction contains ∼ 20-fold higher levels of radioactivity than the extracted Und-P pool in the Δ*abe* strain (Supplemental material, Fig S2B). To this effect, any surplus [^14^C] signals in the LPS fraction derived from the alleviated Und-P would be deemed marginal.

The Group B1 Δ*abe* strains showed 10-fold higher counts for the n-butanol extracts relative to their respective parental strains. Kahne and colleagues reported a similar level of Und-P accumulation (16-fold) when lipid II translocation was blocked in a MurJ variant (35). It should be noted that the kinetics of O-antigen precursor translocation and the subsequent incorporation into LPS are highly efficient in the WT. As such, Und-P pool turnover would occur at a faster pace during the preparation of the WT sample than in strains defective in translocation. Note also that the extracted Und-PP-linked O-antigen pool of the Δ*abe* strain, unlike the WT, would be primarily unpolymerized due to inaccessibility to Wzy polymerase. In such cases, the 10-16-fold sequestration of Und-PP-linked material required for cell death is likely an overestimate. Overall, these observations support that Und-P is available only adequately (or slightly above demand) amounts to accommodate levels required for peptidoglycan and polysaccharide pathways. Recent evidence suggests a feedback control mechanism involving the MraY enzyme, which catalyses the production of lipid I, in that the enzyme possesses a potential lipid II-binding cavity on its extracellular face that is proposed to stall lipid II production in response to excess unincorporated peptidoglycan subunit (36). Hence, regulated flow through shared pathways either by a feedback response as discussed with the MraY example or allocated competitively would be instrumental for bacterial survival, and these elements will need further investigation.

## MATERIALS AND METHODS

### Bacterial strains and growth conditions

The strains and plasmids used in this work are described in Table 1. Nutrient broth (sodium chloride 5 g l^-1^, yeast extract 5 g l^-1^ and bacteriological peptone g l^-1^) with sugar depletion carried out by the aerobic growth of MG1655 as previously described (13) were used for strain maintenance and assessment in this study. Media were supplemented with antibiotics as required at the following concentrations, ampicillin (25 μg ml^-1^), kanamycin (25 μg ml^-1^), and chloramphenicol (12.5 μg ml^-1^).

**Table 1.**
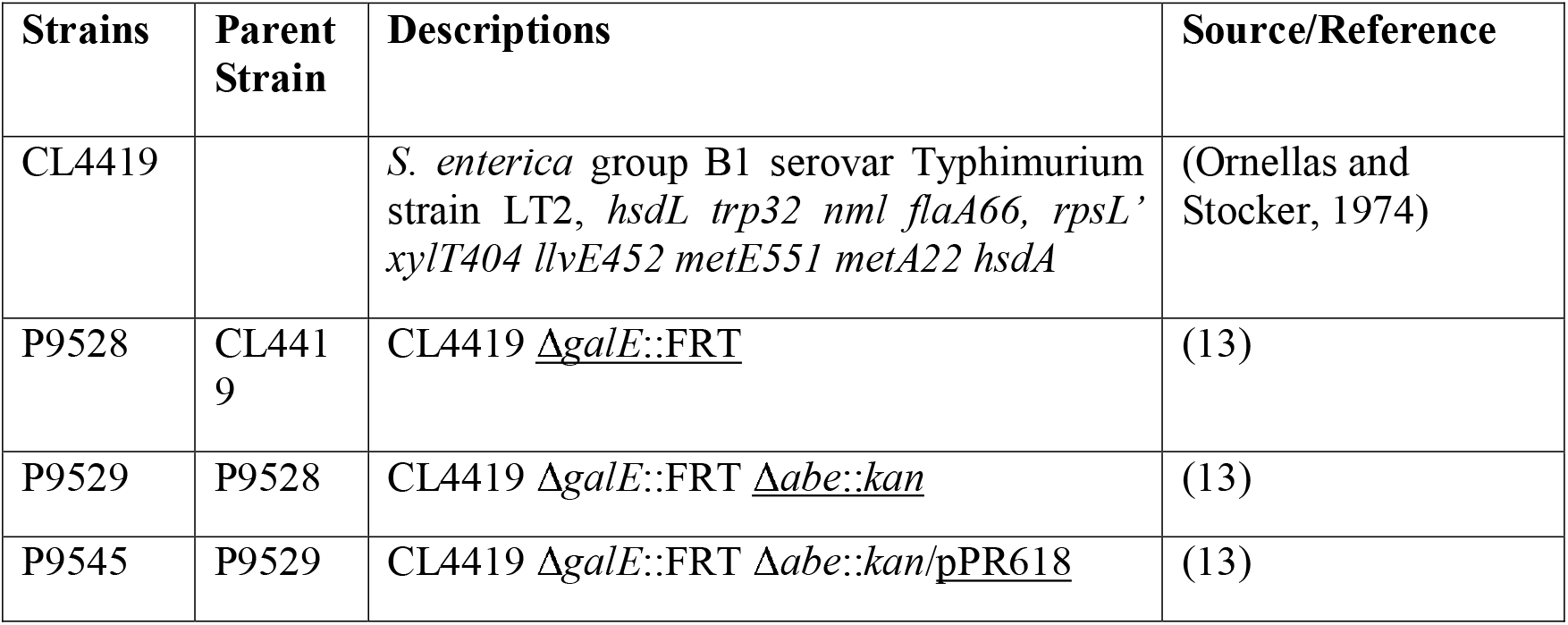
Strain table.

| Strains | Parent Strain | Descriptions | Source/Reference |
| --- | --- | --- | --- |
| CL4419 |  | *S. enterica* group B1 serovar Typhimurium strain LT2, *hsdL trp32 nml flaA66, rpsL' xylT404 llvE452 metE551 metA22 hsdA* | (Ornellas and Stocker, 1974) |
| P9528 | CL4419 | CL4419 <u>$\Delta galE::FRT$</u> | (13) |
| P9529 | P9528 | CL4419 $\Delta galE::FRT$ <u>$\Delta abe::kan$</u> | (13) |
| P9545 | P9529 | CL4419 $\Delta galE::FRT$ $\Delta abe::kan$/<u>pPR618</u> | (13) |

### Assess UDP-galactose epimerase activity

Overnight grown single colonies were streaked onto modified McConkey agar containing 17 g L^-1^ tryptone, 3 g L^-1^ peptone, 10 g L^-1^ D-Gal, 1.5 g sodium deoxycholate, 5 g sodium chloride, 0.03 g neutral red (pH 7.2) and 13 g L^-1^ bacteriological agar. Plates were incubated at 37C for 24 hrs before images were taken.

### [^14^C] D-Gal uptake assay

Overnight broth cultures grown in depleted media were diluted 1:100 into 40mL depleted media. The cultures were incubated at 37C until the mid-log phase. The cultures were then adjusted to equal OD_600_ (∼0.5). [^14^C] D-Gal was mixed with 20% w/v D-Gal to give a specific activity of 35.37 Ci/mol and was then added into the cultures. A total of 250 nCi [^14^C] Gal was added per sample. The cultures were allowed to incubate for another 15 min then rapidly chilled on ice. The cell pellet was then collected and washed in 2 mL 0.85% w/v saline. This sample was loaded into scintillation vials containing 5 mL ACSII scintillant and subjected to quantitative counting in Tri-Cab 2810 TR Liquid Scintillation Analyzer. The scintillation involved counting over 1-minute intervals.

For assessing incorporation into the LPS and n-butanol extractable Und-PP-linked material, equivalent amounts of WT and Δabe culture samples were prepared as above. Each sample was divided into four equal 500 μL lots in 80 mM Tris-acetate (pH 8.0), 10 mM MgCl_2_ and 1 mM EDTA. Two lots were used for hot phenol water LPS extraction as previously described (13). The other two lots were simultaneously added to equivalent volumes of 50 mM pH 8.0 KCl-buffered n-butanol to extract Und-PP-linked material as previously described (25,37). The samples were then vortexed thoroughly to mix. Note that the samples were treated the same until enzymatic activities were deactivated with either phenol or n-butanol. The n-butanol fraction was back-extracted with 80 mM Tris-acetate (pH 8.0), 10 mM MgCl_2_ and 1 mM EDTA twice. One of the two lots used for LPS purification and n-butanol extractions was subjected to sonication. The samples were sonicated in Branson Model 250/450 Stonefire (microtip) for 3 bursts of 8 seconds, with 100% duty cycle and output control calibrated at 3. Experiments were performed in three independent repeats.

## AUTHOR CONTRIBUTION

YH and PRR contributed to project conceptualisation, experimental design, and formal analysis. YH contributed to data collection and drafted the initial manuscript. YH and PRR revised the manuscript.

## REFERENCES

1. TouzÉ, T., and Mengin-Lecreulx, D. (2008) Undecaprenyl Phosphate Synthesis. EcoSal Plus 3

2. Sande, C., and Whitfield, C. (2021) Capsules and Extracellular Polysaccharides in Escherichia coli and Salmonella. EcoSal Plus 9, eESP00332020

3. Hong, Y., Hu, D., Verderosa, A. D., Qin, J., Totsika, M., and Reeves, P. R. (2023) Repeat-unit elongations to produce bacterial complex long polysaccharide chains, an O-antigen perspective. EcoSal Plus

4. Islam, S. T., and Lam, J. S. (2014) Synthesis of bacterial polysaccharides via the Wzx/Wzy-dependent pathway. Canadian Journal of Microbiology 60, 697–716

5. Islam, S. T., and Lam, J. S. (2013) Wzx flippase-mediated membrane translocation of sugar polymer precursors in bacteria. Environ Microbiol 15, 1001–1015

6. Hong, Y., Liu, M. A., and Reeves, P. R. (2018) Progress in Our Understanding of Wzx Flippase for Translocation of Bacterial Membrane Lipid-Linked Oligosaccharide. Journal of Bacteriology 200, e00154–00117

7. Roney, I. J., and Rudner, D. Z. (2023) Two broadly conserved families of polyprenyl-phosphate transporters. Nature 613, 729–734

8. Sit, B., Srisuknimit, V., Bueno, E., Zingl, F. G., Hullahalli, K., Cava, F., and Waldor, M. K. (2023) Undecaprenyl phosphate translocases confer conditional microbial fitness. Nature 613, 721–728

9. Qiao, Y., Lebar, M. D., Schirner, K., Schaefer, K., Tsukamoto, H., Kahne, D., and Walker, S. (2014) Detection of Lipid-Linked Peptidoglycan Precursors by Exploiting an Unexpected Transpeptidase Reaction. Journal of the American Chemical Society 136, 14678–14681

10. Yuasa, R., Levinthal, M., and Nikaido, H. (1969) Biosynthesis of cell wall lipopolysaccharide in mutants of Salmonella V. A mutant of Salmonella typhimurium defective in the synthesis of cytidine diphosphoabequose. Journal of bacteriology 100, 433–444

11. Rick, P. D., and Osborn, M. J. (1972) Isolation of a mutant of Salmonella typhimurium dependent on D-arabinose-5-phosphate for growth and synthesis of 3-deoxy-D-mannooctulosonate. Proceedings of the National Academy of Sciences of the United States of America 69, 3756–3760

12. Rick, P. D., Wolski, S., Barr, K., Ward, S., and Ramsay-Sharer, L. (1988) Accumulation of a lipid-linked intermediate involved in enterobacterial common antigen synthesis in Salmonella typhimurium mutants lacking dTDP-glucose pyrophosphorylase. Journal of Bacteriology 170, 4008–4014

13. Hong, Y., Cunneen, M. M., and Reeves, P. R. (2012) The Wzx translocases for Salmonella enterica O-antigen processing have unexpected serotype specificity. Molecular Microbiology 84, 620–630

14. Liu, M. A., Stent, T. L., Hong, Y., and Reeves, P. R. (2015) Inefficient translocation of a truncated O unit by a Salmonella Wzx affects both O-antigen production and cell growth. FEMS Microbiology Letters 362

15. Hong, Y., and Reeves, P. R. (2016) Model for the Controlled Synthesis of O-Antigen Repeat Units Involving the WaaL Ligase. mSphere 1

16. Xayarath, B., and Yother, J. (2007) Mutations blocking side chain assembly, polymerization, or transport of a Wzy-dependent Streptococcus pneumoniae capsule are lethal in the absence of suppressor mutations and can affect polymer transfer to the cell wall. J Bacteriol 189, 3369–3381

17. Mitchell, A. M., Srikumar, T., and Silhavy, T. J. (2018) Cyclic Enterobacterial Common Antigen Maintains the Outer Membrane Permeability Barrier of Escherichia coli in a Manner Controlled by YhdP. mBio 9

18. Chua, W.-Z., Maiwald, M., Chew Kean, L., Lin Raymond, T.-P., Zheng, S., Sham, L.-T., and Gottesman, S. (2021) High-Throughput Mutagenesis and Cross-Complementation Experiments Reveal Substrate Preference and Critical Residues of the Capsule Transporters in Streptococcus pneumoniae. mBio 0, e02615–02621

19. Maczuga, N., Tran, E. N. H., Qin, J., Morona, R., and El-Naggar, M. Y. (2022) Interdependence of Shigella flexneri O Antigen and Enterobacterial Common Antigen Biosynthetic Pathways. Journal of Bacteriology 0, e00546–00521

20. Jorgenson, M. A., and Young, K. D. (2016) Interrupting Biosynthesis of O Antigen or the Lipopolysaccharide Core Produces Morphological Defects in Escherichia coli by Sequestering Undecaprenyl Phosphate. Journal of Bacteriology 198, 3070–3079

21. Jorgenson, M. A., Kannan, S., Laubacher, M. E., and Young, K. D. (2016) Dead-end intermediates in the enterobacterial common antigen pathway induce morphological defects in Escherichia coli by competing for undecaprenyl phosphate. Molecular Microbiology 100, 1–14

22. Barreteau, H., Magnet, S., El Ghachi, M., Touzé, T., Arthur, M., Mengin-Lecreulx, D., and Blanot, D. (2009) Quantitative high-performance liquid chromatography analysis of the pool levels of undecaprenyl phosphate and its derivatives in bacterial membranes. J Chromatogr B Analyt Technol Biomed Life Sci 877, 213–220

23. Eade, C. R., Wallen, T. W., Gates, C. E., Oliverio, C. L., Scarbrough, B. A., Reid, A. J., Jorgenson, M. A., Young, K. D., and Troutman, J. M. (2021) Making the Enterobacterial Common Antigen Glycan and Measuring Its Substrate Sequestration. ACS Chemical Biology 16, 691–700

24. Hong, Y., and Reeves, P. R. (2014) Diversity of O-Antigen Repeat Unit Structures Can Account for the Substantial Sequence Variation of Wzx Translocases. Journal of Bacteriology 196, 1713–1722

25. Liu, D., Cole, R., and Reeves, P. R. (1996) An O-antigen processing function for Wzx (RfbX): a promising candidate for O-unit flippase. Journal of Bacteriology 178, 2102–2107

26. Rick, P. D., Barr, K., Sankaran, K., Kajimura, J., Rush, J. S., and Waechter, C. J. (2003) Evidence that the wzxE gene of Escherichia coli K-12 encodes a protein involved in the transbilayer movement of a trisaccharide-lipid intermediate in the assembly of Enterobacterial common antigen. Journal of Biological Chemistry 278, 16534

27. Hong, Y., and Reeves, P. R. (2014) Diversity of O-antigen repeat-unit structures can account for the substantial sequence variation in Wzx translocases. Journal of Bacteriology 196, 1713–1722

28. Liu, M. A., Stent, T. L., Hong, Y., and Reeves, P. R. (2015) Inefficient translocation of a truncated O unit by a Salmonella Wzx affects both O-antigen production and cell growth. FEMS Microbiology Letters 362, fnv053

29. Chua, W.-Z., Maiwald, M., Chew, K. L., Lin, R. T.-P., Zheng, S., and Sham, L.-T. (2021) High-Throughput Mutagenesis and Cross-Complementation Experiments Reveal Substrate Preference and Critical Residues of the Capsule Transporters in Streptococcus pneumoniae. mBio 12, e02615–02621

30. Liu, M. A., Morris, P., and Reeves, P. R. (2019) Wzx flippases exhibiting complex O-unit preferences require a new model for Wzx-substrate interactions. Microbiologyopen 8, e00655–e00655

31. Weiner, I., Higuchi, T., Rothfield, L., Saltmarsh-Andrew, M., Osborn, M., and Horecker, B. (1965) Biosynthesis of bacterial lipopolysaccharide. V. Lipid-linked intermediates in the biosynthesis of the O-antigen groups of Salmonella typhimurium. Proceedings of the National Academy of Sciences of the United States of America 54, 228–235

32. Liu, B., Knirel, Y. A., Feng, L., Perepelov, A. V., Senchenkova, S. y. N., Reeves, P. R., and Wang, L. (2014) Structural diversity in Salmonella O antigens and its genetic basis. FEMS microbiology reviews 38, 56–89

33. Liu, B., Furevi, A., Perepelov, A. V., Guo, X., Cao, H., Wang, Q., Reeves, P. R., Knirel, Y. A., Wang, L., and Widmalm, G. (2020) Structure and genetics of Escherichia coli O antigens. FEMS Microbiology Reviews

34. Osborn, M., and Weiner, I. (1968) Biosynthesis of a Bacterial Lipopolysaccharide VI. Mechanism of incorporation of abequose into the O-antigen of Salmonella typhimurium. Journal of Biological Chemistry 243, 2631–2639

35. Qiao, Y., Srisuknimit, V., Rubino, F., Schaefer, K., Ruiz, N., Walker, S., and Kahne, D. (2017) Lipid II overproduction allows direct assay of transpeptidase inhibition by β-lactams. Nature Chemical Biology 13, 793–798

36. Marmont, L. S., Orta, A. K., Corey, R. A., Sychantha, D., Galliano, A. F., Li, Y. E., Baileeves, B. W. A., Greene, N. G., Stansfeld, P. J., William M. Clemons, J., and Bernhardt, T. G. (2023) A feedback control mechanism governs the synthesis of lipid-linked precursors of the bacterial cell wall. bioRxiv, 2023.2008.2001.551478

37. Wang, L., and Reeves, P. R. (1994) Involvement of the galactosyl-1-phosphate transferase encoded by the Salmonella enterica rfbP gene in O-antigen subunit processing. Journal of bacteriology 176, 4348–4356

