## Supplemental material for "Sequestration of dead-end undecaprenyl phosphate-linked oligosaccharide intermediate"

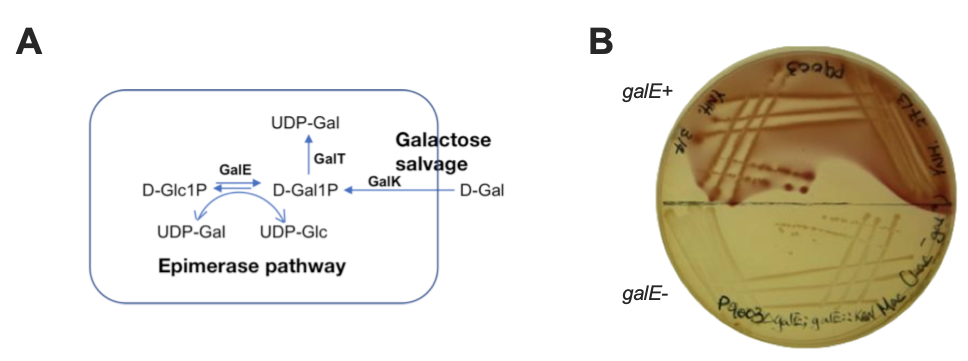

**Fig S1. *Salmonella enterica* vs. Typhimurium ∆*galE* could not metabolise D-Gal. A,** the D-Gal epimerase and salvage pathways; B, D-Gal is not metabolised by *S. enterica* group B1 ∆*galE.*

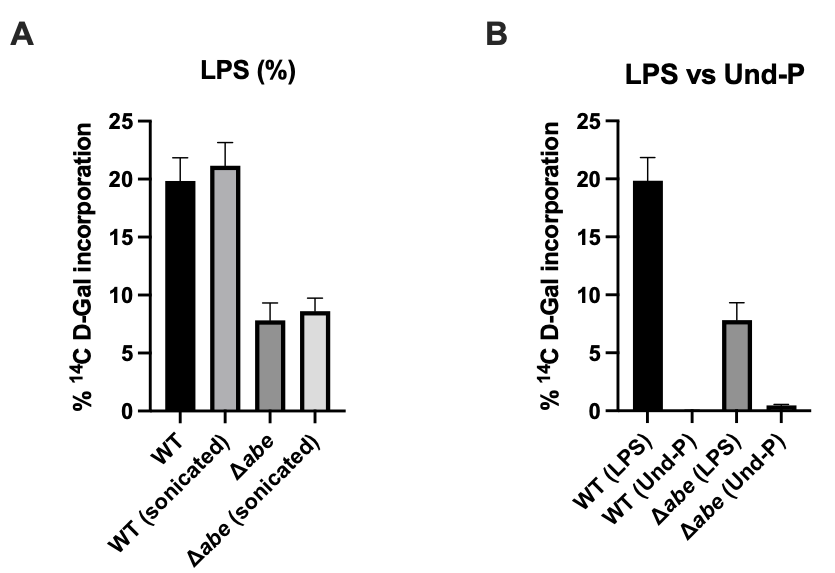

**Fig S2. Effect of sonication-trigged membrane rearrangement on the [^14^C] counts of LPS fraction.** A, Lack of noticeable increment in extracted LPS counts after sonication; B, Relative [^14^C] D-Gal uptake into LPS and Und-P fractions, respectively. Note that the unsonicated LPS and n-butanol extractable pool counts were the same datasets as Fig 2 and Fig 3.
